# Chromosome-associated spot formation by human cytomegalovirus immediate early 1 (IE1) protein

**DOI:** 10.64898/2025.12.18.693715

**Authors:** Mamata Savanagouder, Tejasv Gupta, Martin Messerle, Eva Maria Borst, Thomas F. Schulz, Divya Das, Emma Poole, John Sinclair, Wojciech Zdanowski, Tomasz Waśniewski, Marek Zygmunt, Sławomir Wołczyński, Magdalena Weidner-Glunde

## Abstract

HCMV is associated with severe disease in immunocompromised patients and is a cause of congenital disease. Already more than 30 years ago, IE1 was described to bind to mitotic chromosomes, however the implications of this finding have yet to be determined. HCMV can infect a wide range of cell types and was also detected in different tumours; however, the majority of molecular studies relating to this virus are performed in fibroblasts or monocytic cells. We examined the localization of HCMV IE1 across multiple cell types, including tumour cell lines, to better understand how the protein behaves in these cellular contexts. IE1 is one of the first HCMV genes expressed after lytic infection of the cell and it was observed to distribute evenly over the chromosomes in so-called “chromosome painting” pattern. In leukemia and glioblastoma cells we detected a novel mitotic localization pattern of IE1 - in chromosome-associated spots (CAS), in addition to “painting the chromosomes”, as reported before. We demonstrate that the IE1 core domain mediates CAS localization and that the distribution pattern of IE1 on mitotic chromosomes is influenced by both the cell type being infected and the level of IE1 expression. The novel CAS localization of IE1 suggests a new, possibly cell-type specific function of this protein, distinct from its role in transcriptional regulation.

## Introduction

Infection with human cytomegalovirus (HCMV) is very common and in some countries the prevalence reaches almost 100% of the population (1). Primary HCMV infection in healthy individuals is usually asymptomatic; however, in immunocompromised patients, for example after transplantation or in the context of AIDS, it may cause significant morbidity and mortality (2, 3). HCMV is also the leading infectious cause of congenital disease (4, 5).

After initial acute infection, HCMV persists for the lifetime of the host in latent form, similar to other herpesviruses. The lytic part of the viral infection cycle is associated with active replication and production of new progeny virions. In latency, the virus is persisting in a quiescent mode escaping immune detection, lowering the level of expression of its genes and ensuring persistence of the genome, if in dividing cells. Latency is also defined by the ability of the virus to occasionally reactivate and enter the lytic replication cycle.

The immediate early 1 (IE1) protein of HCMV is one of the first HCMV proteins expressed in the lytic cycle. Together with IE2, IE1 plays a critical role in initiation of early gene expression and viral replication (6, 7). Additionally, it also regulates cellular gene expression by both activation and repression of transcription (reviewed in (8)). IE1 has also been shown to counteract both intrinsic and innate immune responses (reviewed in: (9, 10)).

The IE1 comprises multiple structurally and functionally discrete regions (**Fig.2A**). The N-terminal domain (aa1-24) is predicted to be intrinsically disordered and was shown to contain a nuclear localisation signal (9, 11-13). A core domain, that follows (aa 25-382) is built from 11*α*-helices and adopts a femur-like structural fold (14). This region contributes significantly to stability of the protein, is responsible for IE1 dimerization and contains at least one additional NLS (aa326-342) (9, 11, 12, 14-18). Functionally it was shown to participate in binding to PML (promyelocytic leukemia) protein and in induction of PML nuclear body disruption (9, 11, 12, 16, 18). Adjacent to the core domain is the acidic domain (aa379-475) predicted to be intrinsically disordered, shown to carry signal transducer and activator of transcription (STAT) 2 and 3 binding sites as well as SUMO1 and 3 conjugation sites (13, 19-23); The most C-terminal is the chromosome tethering domain (CTD, aa476-491) shown to mediate IE1 chromatin association and also functioning as a nuclear retention signal (11, 18, 24, 25). The IE1-CTD peptide assumes an extended, V-shaped configuration that includes a short C-terminal α-helix (26).

The functional significance of IE1 association with chromatin is still not fully understood. Thus far, no direct interaction between IE1 and DNA has been demonstrated. A nucleosome binding motif (NBM) found within the CTD (aa 479–488) was shown to mediate the binding of IE1 to histones H2A-H2B (26, 27). IE1 was observed to distribute evenly over the chromosomes in so-called “chromosome painting” pattern (24, 27) and NBM was found to mediate this localization as mutations of these residues led to the release of IE1 from the mitotic chromosomes (27). Although not yet fully elucidated, the association of IE1 with nucleosomes has been linked to modifications in higher-order chromatin structure (26). Binding of histones by IE1 may also be associated with the low nucleosome abundance and the temporal reshaping of nucleosome patterns across viral genomes during productive HCMV infection (28, 29). It has also been suggested that a shorter splice variant of IE1, carrying the CTD domain, namely the IE1x4 protein, tethers HCMV genome to cellular chromatin in a manner analogous to the Kaposi’s sarcoma-associated herpesvirus (KSHV) latency protein LANA (30).

With the goal of studying IE1 localization in tumour cells shown to be HCMV-positive, we uncovered in this study a previously unreported distribution pattern of the full-length IE1 protein, appearing as discrete spots on mitotic chromosomes. We show that the IE1 core domain is necessary and sufficient to mediate this localization. Furthermore, the type of cell that is infected and the level of IE1 protein expression determine its localization pattern on mitotic chromosomes.

## Results

### HCMV IE1 can localize in chromosome-associated spots in mitotic cells

We have performed chromosome spreads in T98G glioblastoma cells. Using this method enabled us to enrich for cells in metaphase and allowed for more detailed analysis of HCMV IE1 localization patterns on mitotic chromosomes, which was the focus of this study. Our experiments showed that in T98G glioblastoma cells, which have previously been successfully used as an HCMV latency model system (31, 32), the HCMV IE1 protein shows two mitotic localization patterns: the previously shown “chromosome painting” and the newly observed chromosome-associated spots (CAS) (**Fig. 1 and S1, S2, S3**). Fig. 1A shows a lower magnification overview of a population of analysed cells enriched in mitosis displaying the two observed patterns. We observed the IE1 CAS both in infected and in transfected cells (**Fig. 1B and C**, respectively). In Fig. 1B and 1C we show representative single cell (single chromosome spread) images of metaphases with the two different IE1 localization patterns identified by us: the “painting” and the CAS. The IE1 CAS staining pattern was detected when using different anti-IE1 antibodies in the context of infection (**Fig. 1B and S1**) and using expression vectors with different tags – myc and GFP (**Fig. 1A, S2 and 1C**, respectively). We have quantified the results of the immunofluorescence staining upon transfection and found that ~39% of mitotic cells showed CAS IE1 pattern (**Fig. 1D**), while in the remaining cells IE1 was painting the chromosomes. In the infected T98G cells it was 24% of mitotic cells that showed the CAS IE1 localization pattern **(Fig. 1D)**. Statistical analysis (Kruskal-Wallis test, with Dunn’s post-test) showed that there is no significant difference between the numbers of cells with CAS pattern in samples transfected with vectors containing different tags (supplementary **Table S1**). The difference between the transfected samples and the infected ones was significant, however this could be related to the higher level of protein expression in infected cells (as will be discussed below). To exclude the possibility that the observed IE1 spotty localization pattern is an artefact of either fixation or chromosome spreading method, we tested three different fixatives: paraformaldehyde, methanol and ethanol and used these on cells processed for regular immunofluorescence staining (without colcemid block and chromosome spread protocol). We were able to detect IE1 CAS in mitotic cells regardless of the fixation method and also without performing chromosome spreads (**Fig. S3**).

**Figure 1.**
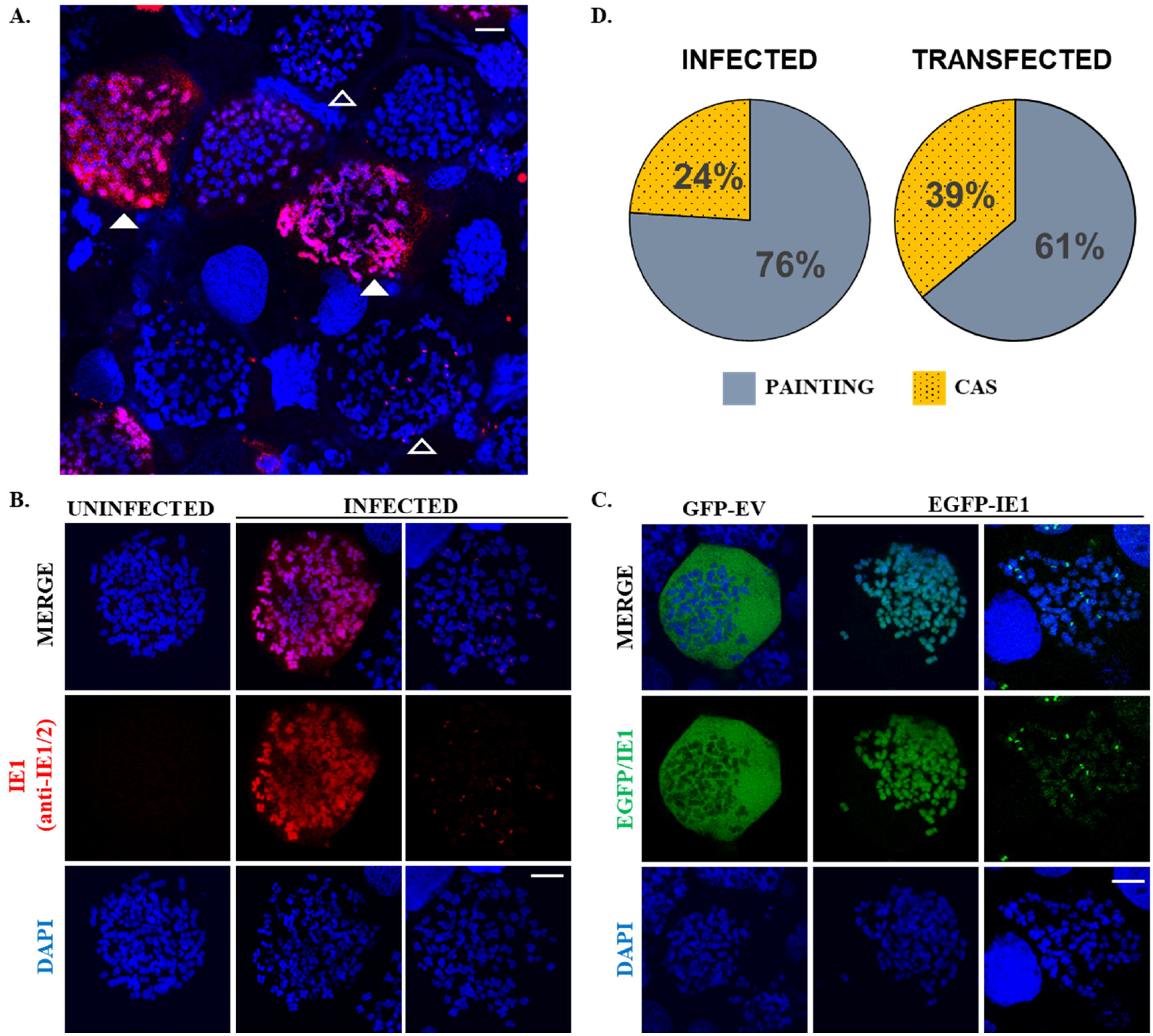
IE1 forms spots on mitotic chromosomes in infected and transfected T98G cells. T98G cells transfected with a plasmid encoding myc-tagged IE1 **(A)**, EGFP-tagged IE1 **(C)** or infected with HCMV **(B)** were arrested in metaphase to perform chromosome spreads. The cells were fixed and stained with anti-myc **(A)**, anti-IE1/2 antibody **(B)** and AF594 conjugated secondary antibody or just stained with DAPI **(C)**. In **(A)** empty arrowheads mark the exemplary cells/chromosome spreads with the CAS pattern and full arrowheads cells with the IE1 “chromosome painting” pattern. In **(B)** and **(C)** in the middle and right panels each image shows a chromosome spread originating from a single cell with either of the two identified IE1 localization patterns. The scale bar represents 10 μm. Quantification of the number of cells with the IE1 painting and CAS localization patterns **(D)** – the full statistical analysis is available in supplementary **Table S1**. Percentages represent the mean from three individual experiments, in each case a total of 100 chromosome spreads of single mitotic cells were counted.

These results suggest that in infected T98G cells IE1 can have an additional novel localization pattern in the form of spots on mitotic chromosomes and that this pattern is formed also in transfected cells, in the absence of other viral proteins or viral genome.

### IE1 core domain is a novel chromatin tethering domain responsible and sufficient for CAS localization pattern

Previous research has identified IE1 CTD and more specifically the NBM within this domain as responsible for mediating interaction of IE1 with the chromosomes (11, 12, 24, 25, 27). The attachment of the IE1 NBM to chromatin has been demonstrated to occur through its interaction with histones, mediating the characteristic “chromosome painting” pattern of IE1. Mutations within the NBM were shown to result in loss of IE1 association with the mitotic chromosomes in fibroblast cells (27). To our knowledge most studies analyzing localization of IE1 on chromosomes so far were performed in fibroblasts (11, 24, 25, 27, 33, 34).

Given the observation of a previously uncharacterized IE1 localization pattern in glioblastoma cells, we sought to examine the behaviour of the IE1 NBM mutant—reported to completely lose chromosome association in fibroblasts — within this cellular context. In glioblastoma cells in part of the cell population, similar to the situation in fibroblasts, IE1 NBM mutant was found off the chromosomes, but in the remaining portion of cells (12%) it was bound to chromosomes in the form of CAS (**Fig. 2B)**. The WT IE1 protein showed two localization patterns: “chromosome painting” and chromosome-associated spots, while the NBM mutant lost the “chromosome painting” pattern, but retained the CAS, suggesting that contrary to the situation in fibroblasts it is still able to bind to chromosomes, but only in the CAS pattern. In addition to IE1 mutant containing alanine substitutions for the most important residues of the nucleosome binding motif, we also confirmed that mutating only the most critical residue M483 of the NBM to alanine gives analogous result (**Fig. 2B**). This suggests that the association of IE1 with the chromosomes as CAS is independent of the CTD/NBM and is mediated by a separate domain.

**Figure 2.**
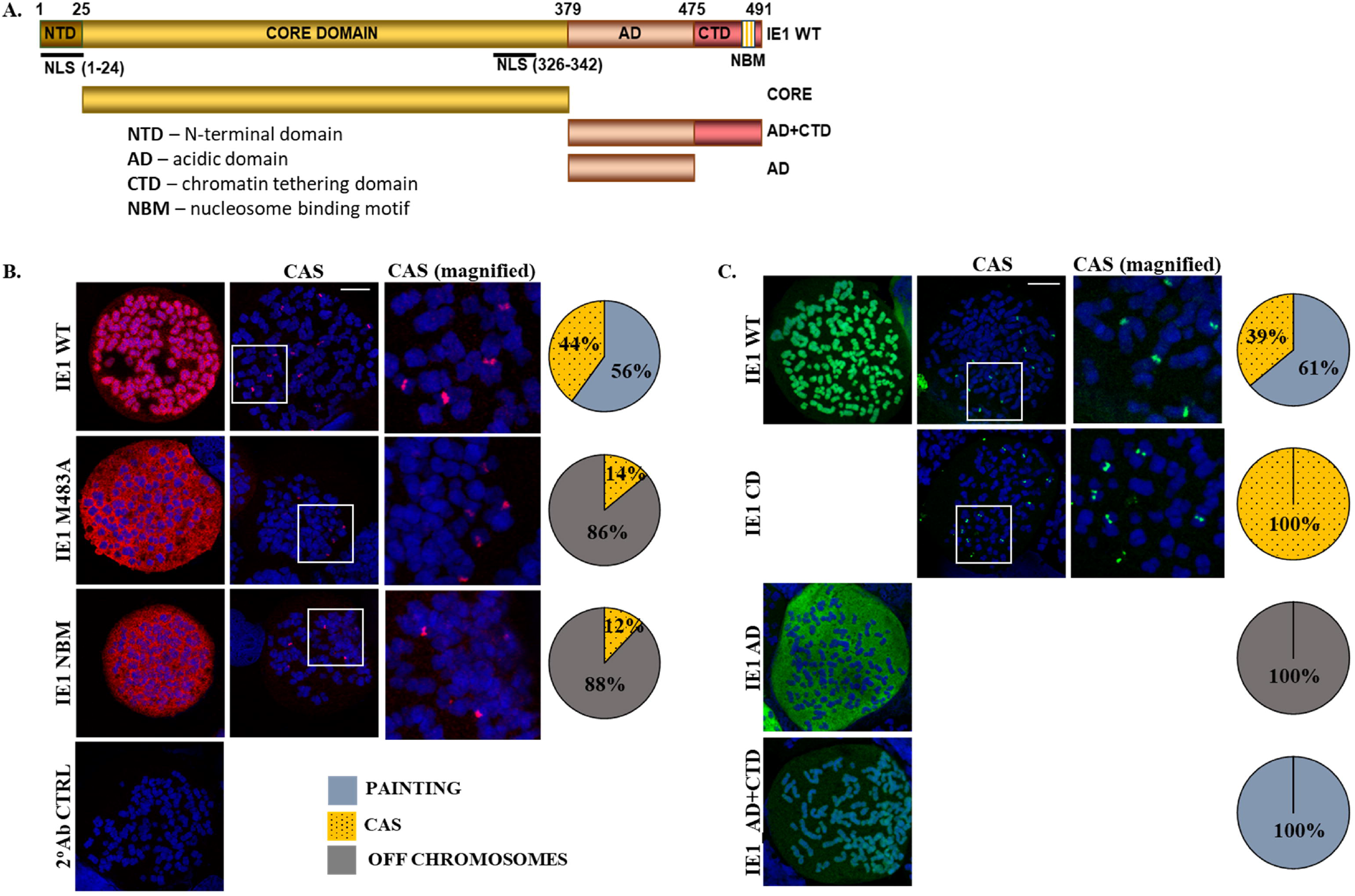
The IE1 core domain is a novel chromatin tethering domain responsible and sufficient for the CAS localization pattern. Constructs used in the study **(A)**. In **(B)** and **(C)** left and middle column, each image shows a chromosome spread originating from a single cell. The third column shows a magnification of the fragment marked on the original CAS image in column two. T98G cells transfected with plasmids encoding myc-IE1, IE1-NBM, and IE1-M483A **(B)** or EGFP-IE1 and IE1 deletion mutants (CD, AD, AD+CTD) **(C)** were arrested in metaphase to perform chromosome spreads. The cells were fixed and stained with anti-myc and AF594 conjugated secondary antibody **(B)** or just stained with DAPI **(C)**. The scale bar represents 10µm. Quantification of IE1 spots, painting and no chromosome association phenotypes is shown for each sample on the right as a pie chart. Results of full statistical analysis are available in supplementary Table S1. Percentages represent the average from three individual experiments, in each case total of 100 chromosome spreads of single mitotic cells were counted.

In order to identify the part of the IE1 that is responsible for the CAS localization pattern we cloned IE1 protein deletion mutants containing different IE1 domains: the core domain (CD), the acidic domain (AD) and AD together with the chromatin tethering domain (AD+CTD) (**Fig. 2A**). We analyzed chromosome spreads of the T98G glioblastoma cells transfected with these IE1 deletion mutants. We found that the isolated IE1 CD localized exclusively in form of chromosome-associated spots and, was therefore necessary and sufficient for the formation of the CAS localization pattern (**Fig. 2C**). The EGFP-IE1-AD mutant localized in an off-chromosomes pattern (100%), and EGFP-IE1-AD+CTD was found to paint the chromosomes (100%) **(Fig.2C)**.

Our data suggest that there are two independent chromatin association domains of IE1: the CTD and the core domain. The CTD is responsible for “the chromosome painting” pattern in all tested types of cells and mediates association with histones. The CD is responsible for the IE1 CAS localization pattern, detected in T98G glioblastoma cells.

### The infected cell type and the IE1 protein expression level influence the IE1 localization pattern

In an attempt to understand what determines the two different localization patterns in one cell population, at one stage of the cell cycle we analysed the mitotic localization of IE1 in different cell types. To this end we performed chromosome spreads in different cell types after infection with a recombinant virus strain (TB40R-ΔRL13-mGFP) (35). We observed the chromosome-associated spot pattern of IE1 only in some transformed cells, namely the monocyte cell line

THP-1, and the glioblastoma cells T98G and U87MG (**Fig. 3)**. We did not detect it in primary human placental fibroblasts (HPF), colorectal cancer cells (HT-29) or neural stem cells (NSC) (**Fig. 3)**. These data suggest that the type of cell infected by HCMV influences the localization of IE1.

**Figure 3.**
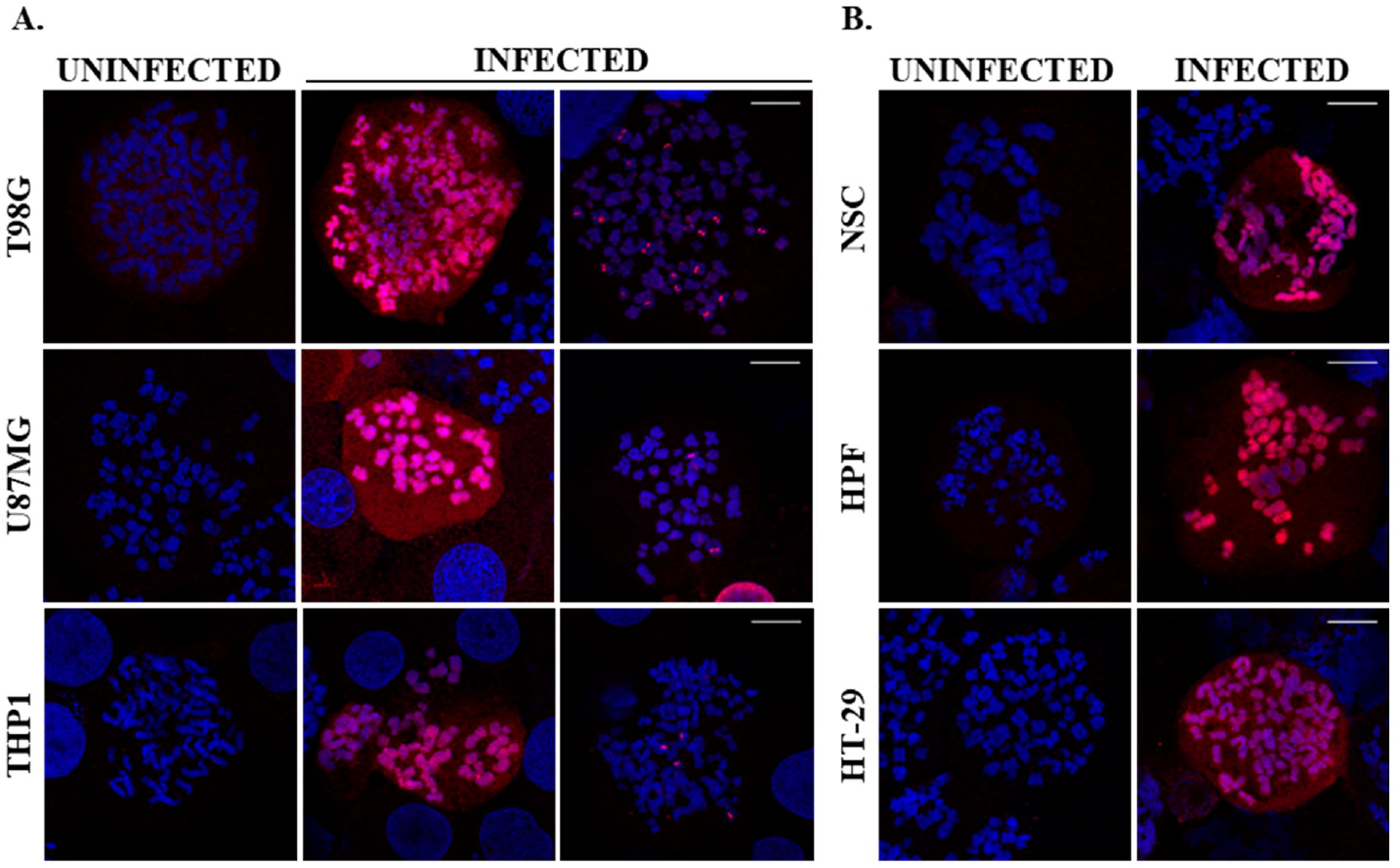
IE1 forms CAS only in some tumor cells. Chromosome spreads of HCMV infected cells were stained with anti-IE1 antibody and AF594 conjugated secondary antibody. Each image shows a chromosome spread originating from a single cell. Cell lines showing IE1 spots on chromosomes in addition to painting pattern were: T98G, U87MG, and THP1 **(A)**, cell lines showing only the IE1 painting pattern were: NSC, HPF, and HT-29 **(B)**. Chromosomes were stained with DAPI. The localization of the stained proteins was analyzed using the 63x objective of a Zeiss LSM800 confocal microscope. The scale bar represents 10 µm. Brightness and contrast were increased for better visualization.

Additionally, we transfected T98G glioblastoma cells with decreasing amounts of IE1 expressing vector and performed chromosome spreads on these cells. We observed that low level of IE1 protein expression (transfection of 0.25 μg IE1 coding plasmid) resulted in domination (75%) of the IE1 CAS localization pattern, while in presence of high IE1 expression levels (transfection of 1.5 μg IE1 coding plasmid) in the majority of cells, the IE1 chromosome painting prevailed (72%) (**Fig. 4 A, B**). We found a negative correlation (Pearson coefficient r = −0.858) between the percentage of cells with IE1 CAS localization pattern and the average level of IE1 expression, as determined by densitometric analysis of the IE1 Western blot signal of samples transfected with decreasing concentrations of the myc-IE1 plasmid (**Fig. 4D**). This indicates that the fraction of cells presenting the IE1 CAS localization pattern is inversely proportional to the level of IE1 protein expression in these cells. These results imply that one of the factors contributing to the localization pattern of IE1 in mitosis is its expression level.

**Figure 4.**
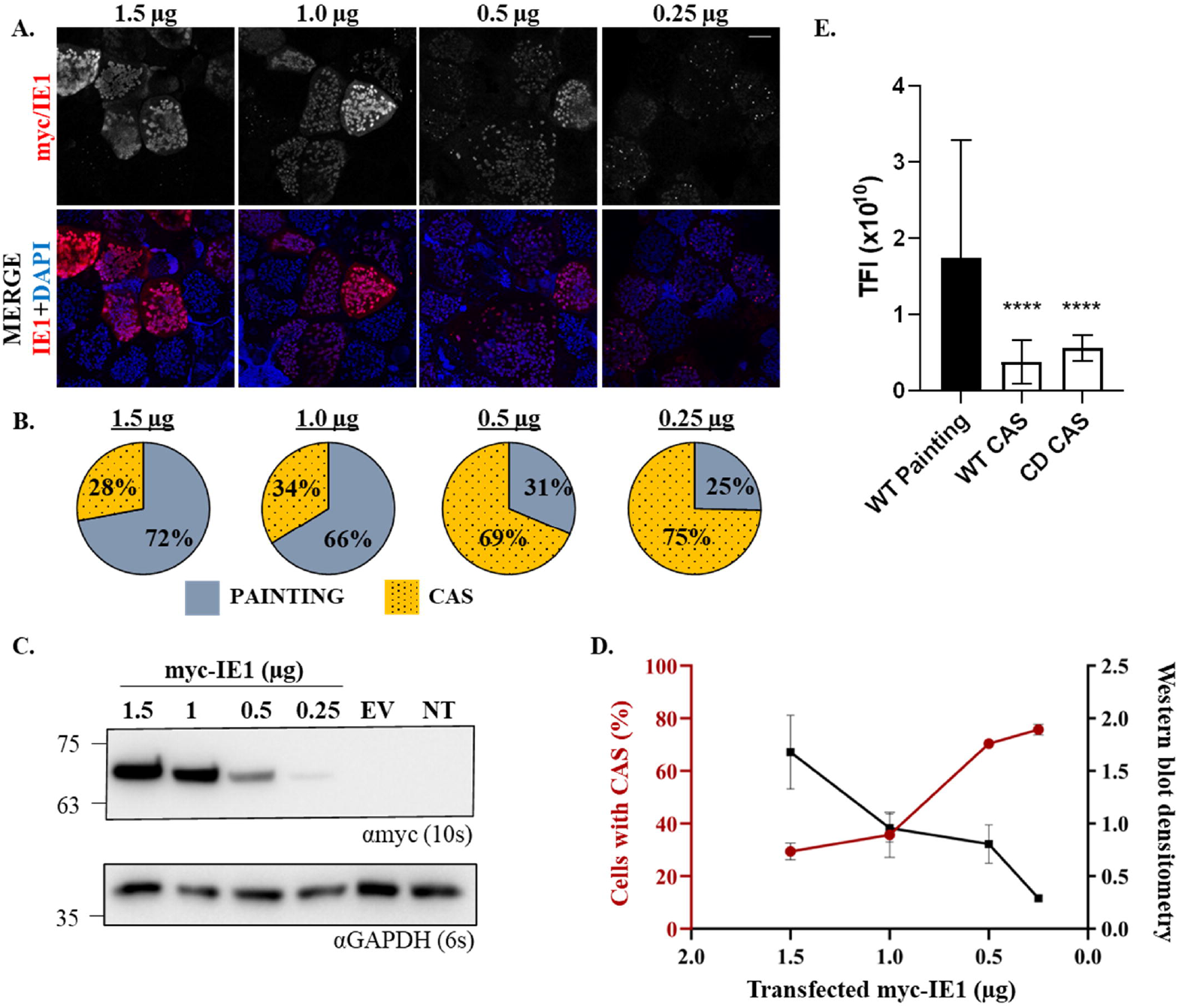
IE1 protein expression level influences its localization pattern in transfected cells. T98G cells transfected with the myc-IE1 WT plasmid were arrested in metaphase to perform chromosome spreads or harvested for immunoblotting. The cells were fixed and stained with anti-myc antibody and AF594 conjugated secondary antibody. Chromosomes were stained with DAPI. The localization of the stained proteins was analyzed using the 40x objective of a Zeiss LSM800 confocal microscope. The scale bar represents 20 µm. The images of cells transfected with decreasing amounts of the myc-IE1 WT plasmid **(A)** and quantification of myc-IE1 painting and CAS localization patterns at the respective concentration **(B)**. Percentages represent the average from three independent experiments. Expression of myc-IE1 analyzed by immunoblotting **(C)**. GAPDH was used as loading control. Antibodies used to detect the proteins are indicated below the blot, the exposure time is indicated in the brackets. EV-empty vector, NT-non transfected. Plot of myc-IE1 protein levels as analyzed by immunoblotting *vs* percentage of cells with CAS localization pattern with decreasing amounts of the used myc-IE1 plasmid **(D)**. Quantification of total fluorescence intensity (TFI) in cells with “chromosome painting” pattern of IE1 WT and CAS pattern of IE1 or IE1 CD **(E)**. The difference in both cases was highly statistically significant (P<0.0001). For each sample minimum 45 cells from 3 experiments were quantified using 3D confocal imaging.

The biggest change in percentage of cells with IE1 CAS pattern was detected following transfection with plasmid amounts between 1.0 μg (34%) and 0.5 μg (69%) (**Fig. 4D**). This suggests that there is a threshold of protein expression level for the switch from IE1 chromosome painting pattern to IE1 CAS. Tukey’s multiple comparison test showed that the difference in percentage of cells with the IE1 CAS localization pattern observed between 1.0 μg and 0.5μg of transfected myc-IE1 plasmid is statistically significant (**Table S1**).

To confirm further the influence of IE1 expression level on its localization pattern in mitosis we quantified the total fluorescence intensity of single cells/chromosome spreads performed in T98G cells in the channel corresponding to the IE1 protein. This analysis showed that the fluorescence intensity is higher in the cells with the “chromosome painting” pattern of IE1 compared to the CAS pattern and that this difference is statistically significant (**Fig. 4E, Table S1**). These results confirm that the expression level of IE1 protein has influence on its localization pattern in mitosis and suggests that changes in IE1 protein expression level lead to changes in localization pattern.

### Limitations of the study

While we demonstrate presence of two distinct patterns of IE1–chromosome association, the underlying molecular mechanism remains incompletely resolved. In particular, we did not perform a comprehensive analysis of the specific protein–protein interactions and chromatin interactions involved in IE1 attachment to chromosomes. Elucidating these interactions requires extensive biochemical and proteomic studies that fall outside the scope of the current work, however it remains a key area for future research. As a result, the present manuscript focuses on describing and comparing the observed localization phenotypes rather than providing a full mechanistic explanation of IE1–chromosome association.

## Discussion

Upon checking the localization pattern of IE1 in T98G glioblastoma cells we observed that in addition to the previously published painting of chromosomes in a substantial fraction of the cells it also forms chromosome-associated spots, which has not been shown before.

Our results suggest that the association of IE1 with the chromosomes in form of chromosome-associated spots is independent of the previously identified chromatin tethering domain and the nucleosome binding motif found within. We identified the core domain as necessary and sufficient for the IE1 CAS localization pattern. However, since the core domain is quite large (355aa), identifying the specific motif responsible for the CAS localization pattern will be an important next step.

In an attempt to understand the mechanism of establishment of the newly discovered IE1 pattern we searched for factors that contribute to its formation. We revealed that the level of expression of the protein influences the localization pattern. Our results suggest that a low level of IE1 expression contributes to a higher percentage of cells showing the IE1 CAS pattern in mitosis, while in presence of a higher level of expression the IE1 “chromosome painting” pattern predominates. The sigmoidal shape of the graph representing the percentage of cells with the CAS localization pattern in relation to the amount of transfected plasmid additionally suggests that there is a threshold, at which the binding mode of IE1 changes (**Fig. 4D**). This implies a potential switch from chromatin-associated spots pattern mediated by the core domain at relatively low level of IE1 expression, to “chromosome painting” pattern mediated by the chromatin tethering domain/ NBM when the IE1 expression level raises sufficiently.

Recently, the level of IE1 expression has been tied to the efficiency of viral entry into cells and the monocytes that received low doses of the virus were more likely to establish a latent state (36). In light of these findings, one could hypothesize that the different mitotic localization patterns of the IE1 protein, determined among others by the expression level of IE1, could be related to different stages of the viral infection cycle. However, whether IE1 CAS corresponds to localization pattern specific to cells that are destined to establish latency is yet to be determined.

Considering that we did not find any published reports describing a chromatin-associated spot-like pattern of IE1, we sought to understand why our observations differed from earlier findings. A careful review of the literature revealed that most prior studies examining IE1 localization on mitotic chromosomes were carried out in fibroblasts (11, 24, 25, 27, 33, 34). Some studies additionally used epithelial cells – Vero (24), HeLa (34), cells with epithelial-like morphology – H1299 lung carcinoma (27) or human embryonic kidney (HEK293) cells (24) - initially considered to be epithelial, but later characterised as neural crest-derived cells. We thus infer that the IE1 mitotic localization is influenced by the experimental cell model and that the observed CAS localization pattern is cell type dependent – so far detected in glioblastoma and monocytic leukaemia cells. Considering that HCMV has a wide cell tropism (37) infecting both somatic tissue cells (fibroblasts, epithelial, endothelial and smooth muscle cells) as well as myeloid cells (monocytes, macrophages, dendritic cells) this finding points to new possible functions of that protein and raises the question whether IE1 might be involved in processes that are cell-type-specific. Further research is required to fully understand this topic.

The punctate association of IE1 with host chromosomes resembles the characteristic and unique localization patterns of established viral episome-maintenance factors, such as KSHV LANA, EBV EBNA1, and HPV E2. These proteins are known to either constitutively form chromosomal foci (E2) or to relocalize into discrete spots upon the presence of viral genomes (LANA, EBNA1) (38-40). Our observations therefore provide a rationale for investigating whether IE1 contributes to HCMV genome maintenance in glioblastoma and leukemia cells.

## Materials and Methods

### Cells and Constructs

T98G (purchased from Sigma, #92090213) and U87MG (purchased from ATCC #ATCC-HTB-14) were cultured in MEM (Cytogen, #04-08500) with non-essential amino acids (Gibco, #11140050), sodium pyruvate (Gibco, #11360070). Human placental fibroblasts (HPF) isolated in our laboratory were cultured in high glucose DMEM with GlutaMAX™ (Gibco), THP1 (kindly provided by Prof. Zygmunt) in RPMI media (Gibco #11875093) and HT-29 (kindly provided by Prof. Wołczyński) in McCoy’ 5A media (Gibco #16600082). All media were supplemented with 10% heat-inactivated FBS (Gibco) and penicillin-streptomycin (Gibco, # 15140122) and the cells were cultured at 37°C with 5% CO2. All cell cultures were regularly confirmed to be mycoplasma negative using MycoSPY from Biontex (#M020-050).

Differentiation of iPSC BIONI010-13 (purchased from Merck/European Bank for induced pluripotent stem cells – EBiSC, #66540023) to neural stem cells (NSC): iPSCs were seeded onto Laminin-111-coated plates (1 μg/cm^2^; Biolamina #LN111) at 1.5×10^5^ cells/well (6-well plate) in differentiation medium 1. From day 0 to 3 10 μM ROCK inhibitor (Sigma #SCM075) was included. On day 4, the medium was switched to differentiation medium 2, and cells were maintained in this medium to sustain the neural stem cell stage. Differentiation Medium 1: DMEM/F12: Neurobasal (1:1; Gibco #11320033, #21103049), 1% N2 (Gibco #17502001), 2% B27 (Gibco #12587001), 10 μM SB431542 (Sigma #616464), 100 ng/ml rhNoggin (Gibco #PHC1506), 300 ng/ml rhSHH-C24II (Sigma #PMC8031), 0.8 μM CHIR99021 (Sigma #SML1046). Differentiation Medium 2: DMEM/F12:Neurobasal (1:1; Gibco #21103049), 0.5% N2 (Gibco #17502001), 1% B27 (Gibco #12587001), 10 μM SB431542 (Sigma #616464), 100 ng/ml rhNoggin (Gibco #PHC1506), 300 ng/ml rhSHH-C24II (Sigma #PMC8031), 0.8 μM CHIR99021(Sigma #SML1046).

A synthetic codon optimized HCMV IE1 gene (Invitrogen) with an N-terminal myc tag containing 5’ HindIII and 3’ BamHI restriction sites was cloned into HindIII and BamHI sites of pcDNA 3.1 (+) to generate a myc-IE1 plasmid. The HCMV IE1 gene with N-terminal EGFP tag was cloned by PCR amplification of the IE1 gene from the myc-IE1 plasmid. The PCR product containing only IE1 sequence (w/o myc tag) was cloned into HindIII- and BamHI-digested pEGFP-C1 to generate the EGFP-IE1 plasmid. Similarly, the IE1 deletion mutants were generated by PCR amplification and cloning into a plasmid containing an N-terminal EGFP fusion. These included the core domain (CD; amino acids 25–378), the acidic domain (AD; aa 379–475) and a combined acidic plus chromatin tethering domain construct (AD+CTD; aa 379–491). Point mutants of IE1—M483A and the mutant M483A_H481A_T485A_R486A (referred to as IE1-NBM) were generated sequentially by site-directed mutagenesis using a previously described PCR protocol (41). Two parallel PCR reactions were performed using the myc-IE1 plasmid as the template, with forward and reverse primers in separate tubes. After amplification, the resulting PCR products were combined, denatured at 95°C, and gradually cooled to 37°C to facilitate reannealing. The mixture was then treated with *DpnI* (NEB) at 37°C for 1 hour to digest the parental plasmid, followed by transformation into TG2 *E. coli* cells. All constructs were sequence verified.

### Virus and virus production

TB40R-ΔRL13-mGFP (35) was grown in HFF cells as described previously (42). Shortly, HFF cells were seeded in T75 flasks at a density of 3×10^6^ cells per flask. On the following day, the cells were infected with HCMV at MOI 0.02 and incubated for 5 h, after which the media was replaced. Once approximately 80-90% of the cells were infected, the supernatants were collected and pre-cleared by centrifugation at 3,500×g for 1 h at 4°C. The virus was concentrated by ultracentrifugation at 103,282×g for 1 h at 4°C in a SW40Ti rotor using a Beckman Optima L-100 XP ultracentrifuge. The pellet containing the viral particles was resuspended in 300 μl of viral stock buffer (50 mM Tris pH 7.8, 12 mM KCl, 5 mM EDTA, 20% heat-inactivated FBS) and stored at −80°C. Titers were determined by a standard plaque assay on HFF cells. Plaques were counted 14 days post-infection following Giemsa staining.

### Infection

T98G infection was carried out as described previously (43). Briefly, serum-starved (78h) T98G cells were plated at 1×10^5^ cells per well of a 6-well plate, allowed to attach for 2h and then infected with TB40R-ΔRL13-mGFP virus at MOI 10. Cells were harvested 4dpi for chromosome spreads.

Neural stem cells (NSCs) were infected on day 3 of differentiation at MOI 3 with centrifugal enhancement (950×g, 30 min, 20°C). Medium was replaced 6h post infection, and cells were harvested 10dpi for chromosome spreads. U87MG cells were seeded at 2×10^5^ cells/well (12-well plate), infected the next day at MOI 2 for 5 h, and then fresh medium was added. Cells were harvested at 2 dpi for chromosome spreads. HT-29 cells were plated at 1×10^5^ cells/well (12-well plate), infected at MOI 1 for 5 h, followed by medium replacement. Cells were harvested at 2 dpi for chromosome spreads. THP-1 cells (1.5×10^6^) were seeded at 0.5×10^6^ cells/ml in 6-well plates and infected at MOI 2 with centrifugal enhancement (450×g, 20min, 20°C). After 5 h incubation, the medium was replaced, and cells were harvested at 2 dpi for chromosome spreads. HPFs were plated at 5×10^5^ cells/well (6-well plate) on coverslips and infected on the next day at MOI 1. Cells were harvested at 2dpi for chromosome spreads.

### Immunofluorescence Assay (IFA)

For interphase staining cells cultured on coverslips (T98G - 3×10^5^ cells per well of a 6-well plate) for transfection (T98G with Lipofectamine 3000 – Invitrogen,) or infection (see above) experiments were washed with PBS and fixed using 4% paraformaldehyde in PBS for 20 min at room temperature (RT). PFA was quenched for 10 min at RT with 125mM glycine and coverslips were washed with PBS. Next, cells were permeabilized with 0.2% Triton X-100 in PBS for 10 minutes at RT and blocked with blocking buffer (5% bovine serum albumin, 0.2% Triton X-100 in PBS) for 30 min at 37°C in a humidifying chamber. Primary antibody staining was performed in blocking buffer for 1h at 37°C in a humidifying chamber, followed by three 5min washes with PBS and incubation with secondary antibody and DAPI (Sigma) in blocking buffer for 1h at 37°C in a humidifying chamber. After additional washes (3 times 5 min with PBS), coverslips were mounted with Mowiol containing DABCO and dried overnight in the dark. Images were captured using a Carl Zeiss LSM800 microscope with a 63x objective, and processed with FIJI software (44). The raw intensity values remained unchanged. Primary antibodies used: rabbit anti-myc (CST, cat #2278, 1:25), mouse anti-IE1/2 (Abcam, cat #ab53495, 1:100), mouse anti-IE1(Santa Cruz, cat. #sc-69834, 1:10). Secondary antibodies used: donkey anti-mouse AF594 conjugated (Jackson Immunolabs cat. #715-585-150, 1:400), donkey anti-rabbit AF594 conjugated (Jackson Immunolabs cat. #711-585-152, 1:200). For the staining of mitotic cells without performing chromosome spreads either 4% PFA was used and the procedure was completed as described above or the coverslips were fixed and permeabilized with ice-cold methanol or 100% ethanol at −20°C for 10 minutes, followed by three PBS washes, after which the procedure was continued with blocking and staining as described above.

### Chromosome Spread

Cells were treated with colcemid (Gibco) using cell type optimized conditions in order to accommodate differences in rates of cell division and in sensitivity to metaphase arrest. The following conditions were used: T98G - 0.2 µg/ml for 6 h, HeLa - 0.1 µg/ml for 12 h, NSC - 0.05µg/ml for 6 h, U87MG and HPF - 0.05µg/ml for 24 h, THP1 - 0.3µg/ml for 6 h, HT-29 - 0.1µg/ml for 12 h. After colcemid incubation, mitotic cells were detached using “mitotic shake off”, centrifuged (10min at 200×g), and the pellet was resuspended in 200 μl of prewarmed 75mM KCl and incubated for 5min at 37°C. Subsequently, cells were span down in a centrifuge (M-DIAGNOSTIC, MPW) with a CYTOSET rotor and adapters (10min at 250×g) onto Superfrost/Plus slides (Thermo Scientific). The cell spot was fixed with 4% paraformaldehyde (20min at RT), quenched with 125 mM glycine (10min at RT), and washed with PBS (3 times 5min at RT). Cells were permeabilized in KCM solution [120 mM KCl, 20 mM NaCl, 10 mM Tris (pH 7.7), 0.1% Triton X-100] 10min at RT, and incubated with primary antibody diluted in KCM solution for 1 h at 37°C in a humidifying chamber. Slides were washed with PBS 3 times 5min at RT and incubated with secondary antibody and DAPI (Sigma) 1h at 37°C in a humidifying chamber. Following washes with PBS (3 times 5min at RT) slides were mounted with Mowiol containing DABCO (Sigma), dried, and imaged using a Carl Zeiss LSM800 microscope 63x objective at 2x digital zoom. Images were processed using FIJI software (44). The raw intensity values remained unchanged. Primary and secondary antibodies used were as described for the IFA assay.

The quantification of percentage of cells/chromosome spreads with IE1 CAS pattern presented in Figs. 1D, 2B,C and 4D was performed by counting number of single cell chromosome spreads with each of the patterns observed for particular sample and presenting it as a percentage of total number of counted cells. A minimum of 100 cells per experiment in 3 independent experiments was counted.

Total fluorescence intensity was quantified in 3D (Fig. 4E). We acquired confocal z-stacks spanning the full cell volume using Leica TCS SP8 X with white laser. The images were acquired in 488nm channel, corresponding to IE1 signal. Pixel size was set according to Nyquist to capture the highest possible resolution. Obtained images were analysed using Imaris software (Oxford Instruments/Bitplane). For each sample minimum 45 cells from 3 experiments were quantified.

### Western Blot

Cell lysates were prepared from 3×10^5^ T98G cells transfected using Lipofectamine 3000 and harvested 48h post transfection. Cells were suspended in 300µl of lysis buffer [50 mM Tris (pH 7.6), 100 mM NaCl, 0.5 mM EDTA, 1% glycerol, and 0.2% IGEPAL], supplemented with protease inhibitors [1.5µM aprotinin (Applichem, #A2132), 10µM leupeptin (Applichem, #A2183), 100µM phenylmethylsulfonyl fluoride (Applichem, #A0999), 1µM benzamidine (Sigma, #434760), and 1.46µM pepstatin A (Applichem, #A2205)], phosphatase inhibitors [1mM sodium fluoride (Sigma, #201154) and 1mM sodium orthovanadate (Sigma, #S6508)], and a de-sumoylating enzyme inhibitor [250µM N-ethylmaleimide (Sigma, #E1271)]. The lysates were sonicated, cleared by centrifugation and the total protein concentration was adjusted based on the Quick Start™ Bradford Protein Assay Kit (Bio-Rad #5000201). Proteins were separated on an SDS polyacrylamide gel, transferred to a nitrocellulose membrane and detected using following antibodies: rabbit anti-myc (CST, cat #2278, 1:1000), rabbit anti-GAPDH (CST, cat/# 2118S). Secondary antibodies used: HRP-conjugated goat anti-rabbit IgG (1:2000, Dako #P0448). Signal was visualized using SuperSignal™ West Pico PLUS Chemiluminescent Substrate (Thermo Fisher Scientific #34577) and the ChemiDoc Touch Imaging System (Bio-Rad).

## Statistical Analysis

All statistical analyses were performed using the GraphPad Prism 10 software. The results are presented in supplementary Table S1.

Kruskal-Wallis test with Dunn’s multiple comparison test was performed to compare percentages of chromosome spreads with IE1 CAS pattern for the following groups: infected T98G cells with cells transfected with myc-IE and with EGFP-IE1. The only statistically significant difference detected in this test was between myc-IE and the infected sample.

Kruskal-Wallis test with Dunn’s multiple comparison test was also performed to compare percentages of chromosome spreads with IE1 CAS pattern between IE1 WT sample and both the M483A and NBM mutants (Fig.2B) as well as the deletion mutants (Fig.2C). No statistical difference was detected between the IE1 WT and these samples, although this is likely due to low sample size.

One-way ANOVA with Tukey’s post-test was performed to compare the percentages of chromosome spreads with IE1 CAS pattern between samples transfected with decreasing amounts (1.5μg, 1.0 μg, 0.5 μg, 0.25 μg) of the myc-IE1WT plasmid (Fig.4B).

Pearson correlation coefficient (r) was determined for percentages of cells with CAS localization pattern vs IE1 band intensity in a western blot across different amounts of transfected myc-IE1 (0.25, 0.5, 1.0, and 1.5μg) (Fig.4D). Homogeneity of variance was confirmed using the Brown-Forsythe test and normality was confirmed using Shapiro-Wilk test. Kruskal-Wallis test with Dunn’s multiple comparison test was performed to compare total fluorescence intensity (TFI) per cell/chromosome spread between cells presenting IE1 painting pattern and cells with CAS (Fig.4E). The difference between IE1 WT painting pattern and both the CAS patterns (IE1 WT and IE1 CD) was highly statistically significant.

The studies involving human placental fibroblasts were approved by Bioethical Committe at the Warminsko-Mazurskie Chamber of Physicians in Olsztyn, Poland (Warminsko-Mazurska Izba Lekarska w Olsztynie) decision nr 17/2019/VII. The studies were conducted in accordance with the local legislation and institutional requirements.

## Acknowledgements

This work was supported by the National Science Centre in Poland grant no. 2017/26/E/NZ6/01124 to MWG

We thank dr Rudolf Bauerfeind and Oliver Terwolbeck from the Research Core Unit for Laser Microscopy, Hannover Medical School for the help at the stage of quantification of fluorescence data and MSc Aravind Ravi for help with data analysis.

## Author contributions

performed research – MS, TG, MWG, EMB, DD, EP; analysed data – MS, MWG, TG; conceived and designed experiments – MWG, MM, MS; provided reagents – TFS, JS, EP, MM, EMB, MZ, SW, WZ, TW; wrote the manuscript – MWG, TG, edited manuscript – MS, TG, EMB, TFS, DD, EP, JS, WZ, TW, MZ, SW, MWG. All authors have read and agreed to the published version of the manuscript.

## Figure legends

**Figure S1. IE1 chromosome associated spots (CAS) are present in HCMV infected T98G cells**. The images show representative chromosome spreads of single cells with either of the two identified IE1 localization patterns. Infected or uninfected T98G cells were arrested in metaphase to perform chromosome spreads. The cells were fixed and stained with anti-IE1x4 antibody and AF594 conjugated secondary antibody. Chromosomes were stained with DAPI. The scale bar, 10 μm.

**Figure S2. IE1 forms chromosome associated spots (CAS) in a proportion of transfected T98G cells**. The images show representative chromosome spreads of single cells with either of the two identified IE1 localization patterns. T98G cells transfected with the myc-tagged IE1 plasmid were arrested in metaphase to perform chromosome spreads. The cells were fixed and stained with anti-myc antibody and AF594 conjugated secondary antibody. Chromosomes were stained with DAPI. The scale bar represents 10 µm.

**Figure S3. IE1 CAS can be observed in mitotic cells fixed with different methods**. T98G cells transfected with the myc-IE1 plasmid were fixed with paraformaldehyde **(A)**, methanol **(B)**, or ethanol **(C)**. The cells were stained with anti-myc antibody and AF594 conjugated secondary antibody. Nuclei were stained with DAPI. The scale bar represents 10µm. No arresting agent was used; mitotic cells were selected from the whole population of unsynchronized cells.

