## Supplementary figures and images for "Chromosome-associated spot formation by human cytomegalovirus immediate early 1 (IE1) protein"

### Fig.S1

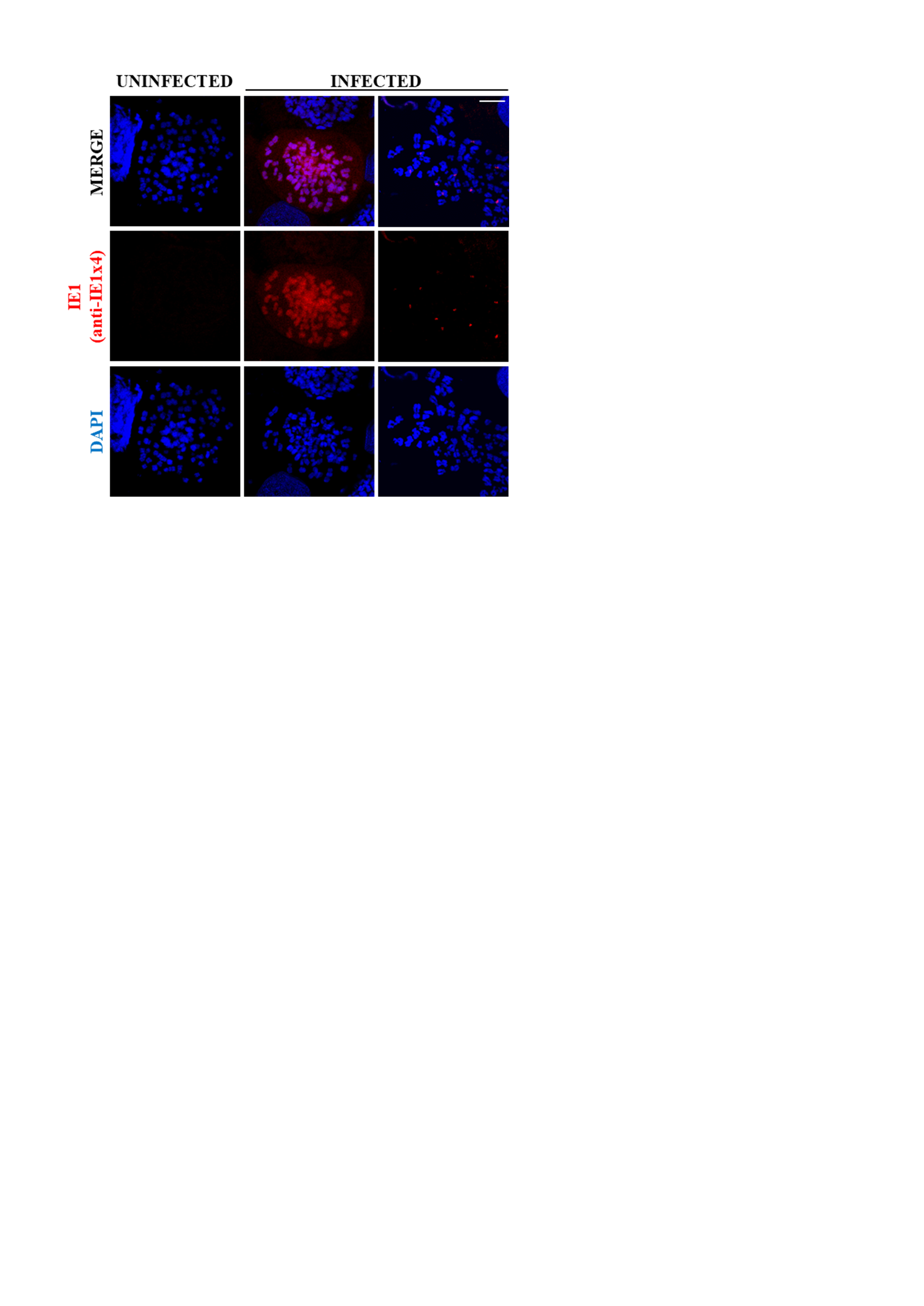

### Fig.S2

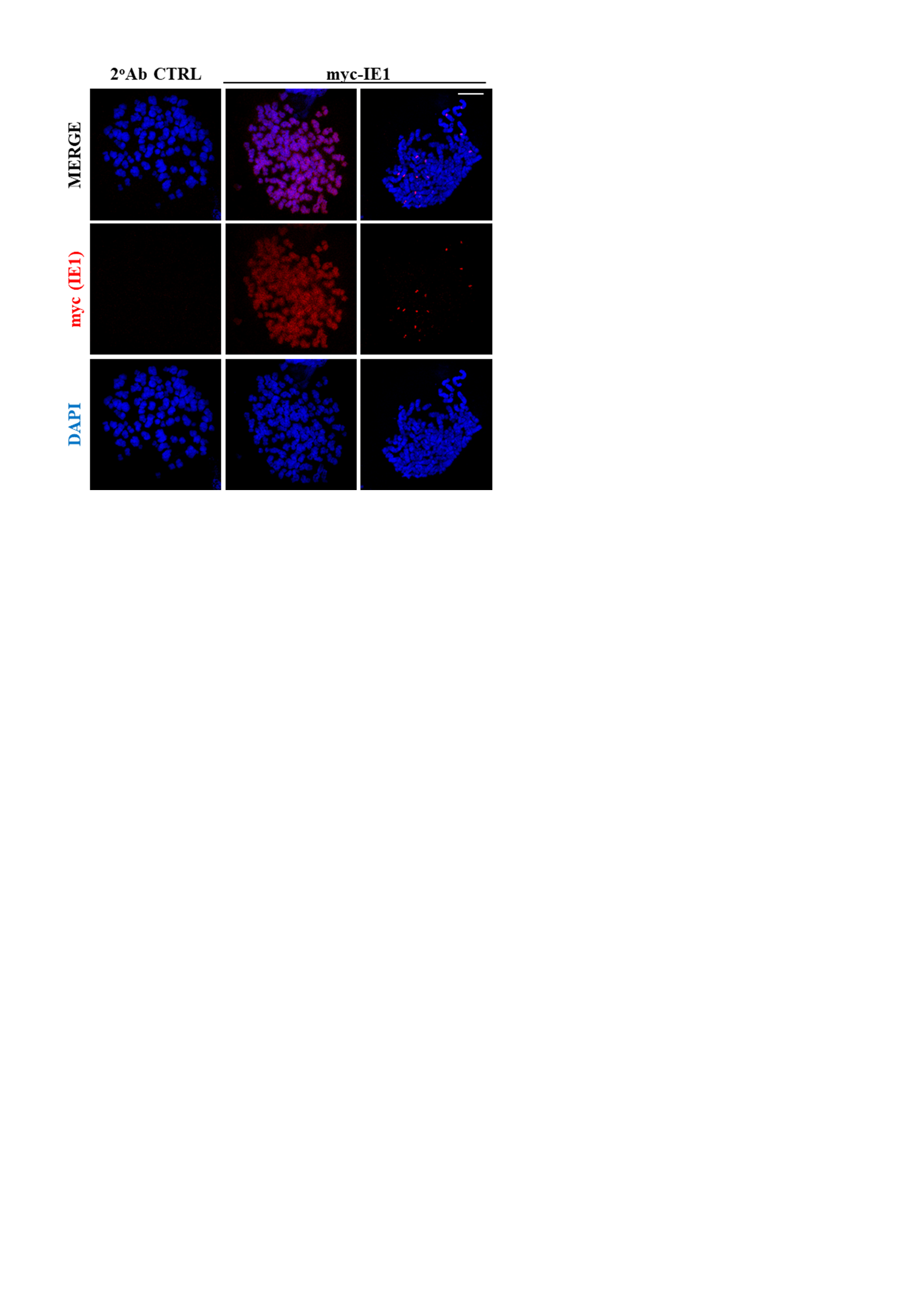

### Fig.S3

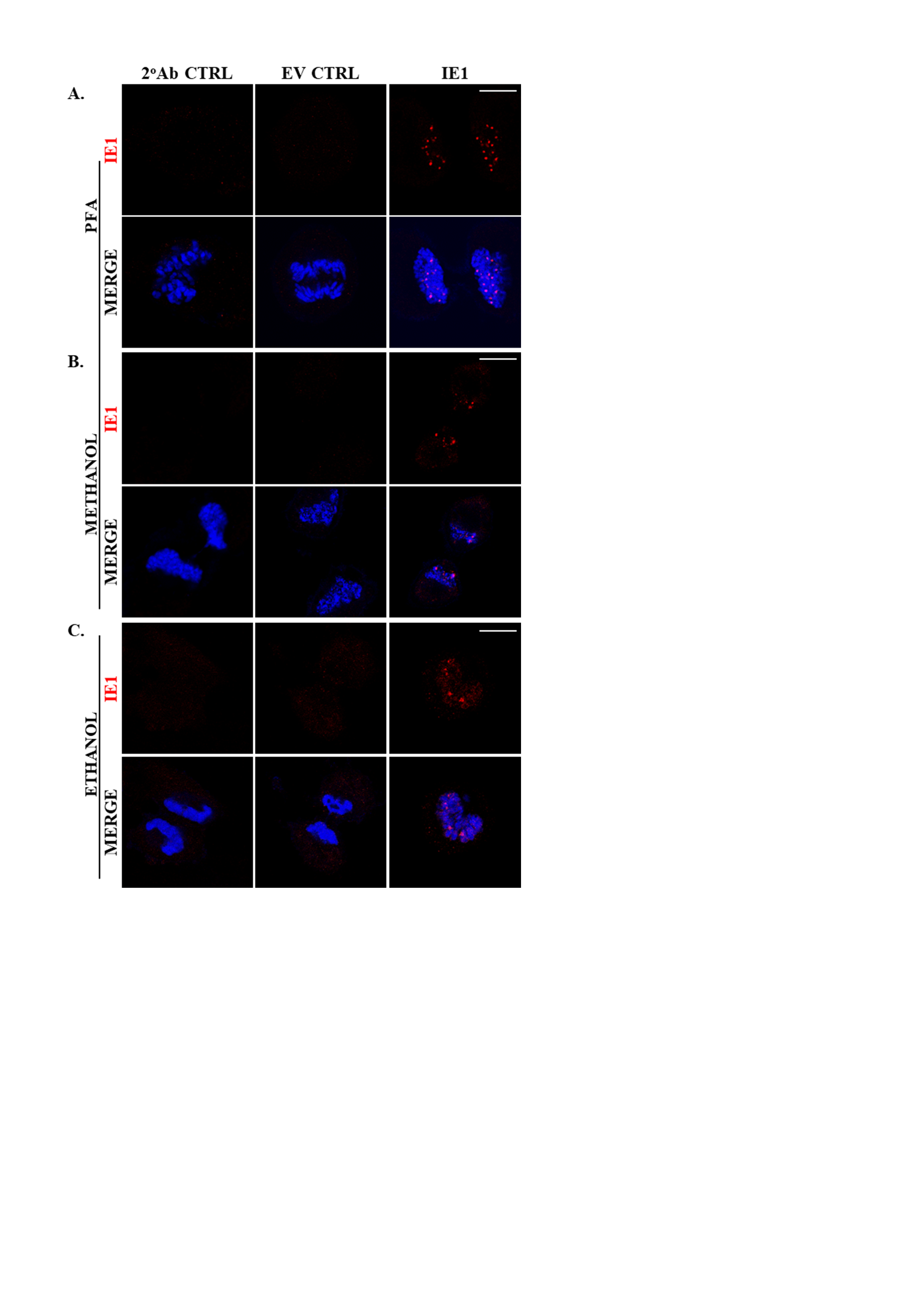
