## Supplementary material for "Chromosome-associated spot formation by human cytomegalovirus immediate early 1 (IE1) protein": Table S1

**Fig.1 D** **Comparison of % of chromosme spreads with CAS in transfected and infected cells**

| | c-myc-IE1 | EGFP-IE1 | D4 INF |
|---|---|---|---|
| | 49 | 36 | 22 |
| | 47 | 39 | 27 |
| | 39 | 42 | 25 |
| **Mean** | **45** | **39** | **25** |
| **STDEV** | **5.3** | **3.0** | **2.5** |

**Kruskal-Wallis test with Dunn's post test**

| **Dunn's multiple comparisons test** | Significant? | Summary | Adjusted P Value |
|---|---|---|---|
| myc-IE1 vs. EGFP-IE1 | No | ns | >0.9999 |
| myc-IE1 vs. D4 INF | Yes | * | 0.0405 |
| EGFP-IE1 vs. D4 INF | No | ns | 0.348 |

**Fig. 2B** **Comparison of % of chromosme spreads with CAS between IE1 WT and M**

| | IE1WT | IE1 M483A | IE1 NBM |
|---|---|---|---|
| | 49 | 18 | 14 |
| | 47 | 16 | 16 |
| | 39 | 11 | 10 |
| **Mean** | **45** | **15** | **13** |
| **STDEV** | **5.3** | **3.6** | **3.1** |

**Kruskal-Wallis test for single pattern (CAS) with Dunn's post test**

| **Dunn's multiple comparisons test** | Significant? | Summary | Adjusted P Value |
|---|---|---|---|
| IE1WT vs. IE1M483A | No | ns | 0.2555 |
| IE1WT vs. IE1NBM | No | ns | 0.061 |
| IE1M483A vs. IE1NBM | No | ns | >0.9999 |

**Fig. 2C** **Comparison of % of chromosme spreads with CAS between IE1 WT and CD/AD/AD+CTD deletion mutants**

| | WT | CD | AD | AD+CTD |
|---|---|---|---|---|
| | 36 | 102 | 0 | 0 |
| | 39 | 100 | 0 | 0 |
| | 42 | 103 | 0 | 0 |
| **Mean** | **39** | **102** | **0** | **0** |
| **STDEV** | **3.0** | **1.5** | **0.0** | **0** |

**Kruskal-Wallis test for single pattern (CAS) with Dunn's post test**

| **Dunn's multiple comparisons test** | Significant? | Summary | Adjusted P Value |
|---|---|---|---|
| WT vs. CD | No | ns | >0.9999 |
| WT vs. AD | No | ns | 0.6165 |
| WT vs. AD+CTD | No | ns | 0.6165 |
| CD vs. AD | Yes | * | 0.0392 |
| CD vs. AD+CTD | Yes | * | 0.0392 |
| AD vs. AD+CTD | No | ns | >0.9999 |

**Fig. 4B** **Comparison of % of chromosme spreads with CAS between samples with IE1 WT transfection in decreasing amounts**

| **Plasmid amount (ug)** | **1.5** | **1.0** | **0.5** | **0.3** |
|---|---|---|---|---|
| **CAS** | 27 | 28 | 69 | 74 |
| | 28 | 34 | 72 | 75 |
| | 33 | 45 | 70 | 78 |
| **Mean** | 29 | 35 | 70 | 76 |
| **STDEV** | 3.2 | 8.6 | 1.5 | 2.1 |

**One Way ANOVA test for single pattern (CAS) with Tukey's post test**

| **Brown-Forsythe test** | |
|---|---|
| F (DFn, DFd) | 1.366 (3, 8) |
| P value | 0.3209 |
| P value summary | ns |
| Are SDs significantly different (P < 0.05) | No |

| **Shapiro-Wilk (W)** | Statistics | P value | Passed normality test ( P value summary |
|---|---|---|---|

| | | | |
|---|---|---|---|
| 0.9085 | 0.204 | Yes | ns |

| **Tukey's multiple comparisons test** | Below threshold? | Summary | Adjusted P Value |
|---|---|---|---|
| 1.5 vs. 1.0 | No | ns | 0.4187 |
| 1.5 vs. 0.5 | Yes | **** | <0.0001 |
| 1.5 vs. 0.25 | Yes | **** | <0.0001 |
| 1.0 vs. 0.5 | Yes | **** | <0.0001 |
| 1.0 vs. 0.25 | Yes | **** | <0.0001 |
| 0.5 vs. 0.25 | No | ns | 0.5508 |

**Fig. 4F** **Comparison of total fluorescence intensity between painting and CAS patterns**

**Kruskal-Wallis test with Dunn's post test**

| **Dunn's multiple comparisons test** | Significant? | Summary | Adjusted P Value |
|---|---|---|---|
| WT Painting vs. WT CAS | Yes | **** | <0.0001 |
| WT Painting vs. CD CAS | Yes | **** | <0.0001 |
| WT CAS vs. CD CAS | Yes | * | 0.0352 |
